# Supplementary motor area in speech initiation: a large-scale intracranial EEG evaluation of stereotyped word articulation

**DOI:** 10.1101/2023.04.04.535557

**Authors:** Latané Bullock, Kiefer J Forseth, Oscar Woolnough, Patrick S Rollo, Nitin Tandon

## Abstract

Speech production is known to engage a distributed network. The brain regions responsible for the initiation of articulation are unclear, and they would be expected to coordinate a distributed network. Using direct intracranial recordings in a large cohort, during stereotyped speech production to eliminate confounds of phonological and semantic complexity, we performed a comprehensive analysis of cortical sites engaged in speech initiation. We found that the supplementary motor area (SMA) was the earliest speech motor region to be active prior to speech onset and was active almost exclusively prior to articulation. Broadband gamma activity in the SMA was predictive of the response time of word production, predicting response time from 250 ms after stimulus onset. Neural activity in SMA began at a highly predictable time after stimulus onset and extended until speech onset. SMA activity *offset* coincided with ventral premotor cortex and primary motor activity *onset*. This suggests that the SMA may coordinate the concerted activation of motor execution cortex. Our results support the idea that SMA is a key node in the speech initiation network.

**Significance statement:** Producing speech requires coordination across multiple brain regions. One cortical region - the supplementary motor area (SMA) - has long been a candidate region to coordinate across other regions to initiate speech production. In this study, we used human intracranial recordings from patients with epilepsy to investigate the role of the SMA in initiating speech. In a picture-naming task, subjects repeated the word “scrambled” scores of times; using this condition to eliminate any linguistic confounds, we found that the SMA was consistently one of the earliest regions to activate during word production. We also uncovered the SMA’s temporally precise tuning to speech onset at the single-trial level.

## Introduction

Initiation of voluntary movement requires coordinated activity between higher-order cognition regions (specifying movement goals) and lower-level sensorimotor regions (specifying sensory feedback and motor plans). The supplementary motor area (SMA) has long been proposed as a critical node in coordinating these processes across many voluntary movements, including propositional speech in humans (Alario et al., 2006; Jonas, 1981; Krainik et al., 2003; Penfield & Welch, 1951). Its stimulation can result in speech arrest (Lu et al., 2021), and SMA lesions may result in impaired propositional speech (Krainik et al., 2003; Laplane et al., 1977). Yet, it’s actual role in speech production remains poorly characterized due to temporal constraints (in fMRI), signal quality in medial areas (as in MEG; e.g., Goldenholz et al., 2008), or sparse coverage (intracranial recordings; e.g., Castellucci et al., 2022). Data from non-human primates reveal that SMA and pre-SMA play an important role in initiating (Kermadi, Y. Liu, A. Tempini E.M. Ro, 1997) and sequencing (Shima & Tanji, 2000; Tanji & Shima, 1994) limb movements. In humans, the pre-SMA and SMA have robust structural connectivity via the frontal aslant tract to ventral premotor and sensorimotor speech cortices, concordant with a putative role for the SMA initiating a ‘go’ signal or ‘igniting’ speech motor output (Catani et al., 2012; Dick et al., 2019). fMRI studies have provided some evidence that the caudal aspect of the pre-SMA/SMA complex controls motor output (Alario et al., 2006; Bohland & Guenther, 2006; Brendel et al., 2010). Some current models of speech production hypothesize that the cascade of speech motor areas begins with coordination between basal ganglia and mesiofrontal cortex (Guenther, 2016), but direct, temporally-precise evidence for this in humans is lacking.

Human intracranial recordings have been key to elucidating the neurobiology of speech articulators (Chartier et al., 2018; Conant et al., 2014; Mugler et al., 2018) and speech initiation (Castellucci et al., 2022; Conner et al., 2019; Forseth et al., 2018; Woolnough et al., 2022), but these studies have mostly been limited to analyses of lateral frontal and Rolandic gyral cortex. We utilized fine-grained intracranial EEG recordings to analyze the activity of the SMA relative to the activation of the speech motor network via large-scale intracranial recordings (n = 115 patients, 125 implants) in a picture-naming task (Forseth et al., 2021). Penetrating depth electrodes covered the broad articulation network, including the medial frontal cortex (i.e., preSMA and SMA). Further, we achieve unprecedented temporal precision by analyzing the repetition of a single stereotyped word - ‘scrambled’ – that participants produced scores of times in response to a jumbled image (8,078 total trials). This enabled the investigation of trial-by-trial variations in response time, without the confound of varying phonological or semantic features.

## Methods

### PARTICIPANTS

After obtaining written informed consent, we enrolled 115 patients (54 men, 61 women; mean age 33 ± 10 years; mean IQ 96 ± 14, 10 left-handed) undergoing evaluation of intractable epilepsy with intracranial electrodes (Forseth et al., 2021). Eight patients were each implanted twice each and one patient was implanted three times, for a total of 125 subject sessions. The study design was approved by the committee for the protection of human subjects at The University of Texas Health Science Center as Protocol Number HSC-MS-06-0385. Nine additional patients recorded from were excluded from analysis as they were determined to be right hemisphere dominant for language.

### EXPERIMENTAL PARADIGM

Patients engaged in a picture naming task (Conner et al., 2014; Forseth et al., 2018, 2021; Woolnough et al., 2019). They were instructed to articulate the name of common objects depicted by line drawings (Snodgrass & Vanderwart, 1980) as quickly and accurately as possible. A control condition was intermixed consisting of the same images with pixel blocks randomly rotated; for these trials, patients were instructed to respond with “scrambled.” Each visual stimulus was displayed for 2 seconds on a 15-in LCD screen positioned at eye level, with an interstimulus interval of 3 seconds. A minimum of 120 (mean 298) visual stimuli were presented to each patient using stimulus presentation software (Python v2.7). 66 ± 18 (mean ± SD) scrambled trials were presented per patient. In total, the word ‘scrambled’ was produced 8,078 times across the cohort. Mean accuracy was >90% in all patients.

### ELECTRODE IMPLANTATION AND DATA RECORDING

Subdural grid electrodes (SDEs; n = 39 implants) – subdural platinum-iridium electrodes embedded in a silicone elastomer sheet (PMT Corporation, top-hat design; 3-mm diameter cortical contact) – were surgically implanted via a craniotomy (Tandon & Luders, 2008). Electrocorticography recordings were performed at least 2 days after the craniotomy. Stereo-electroencephalography (sEEG; n = 86 implants) probes contained platinum-iridium electrode contacts (PMT Corporation; 0.8-mm diameter, 2.0-mm length cylinders; separated from adjacent contacts by 1.5 to 2.43 mm), and were implanted using ROSA, with stereotactic skull screws registered to both a computed tomographic angiogram and an anatomical MRI (Gonzalez-Martinez et al., 2014; González-Martínez et al., 2016; Rollo et al., 2020; Tandon et al., 2019). There were 8 to 16 contacts along each depth probe, and each patient had multiple (12 to 20) probes implanted. Intracranial EEG data were collected with a sampling rate of 1 kHz and bandwidth of 0.15 to 300 Hz using Neurofax (Nihon Kohden) or with a sampling rate of 2 kHz and bandwidth of 0.1 to 700 Hz using NeuroPort NSP (Blackrock Microsystems). Continuous audio recordings were performed with both an omnidirectional microphone (Audio Technica U841A, 30 to 20,000 Hz response, 73 dB SNR) placed adjacent to the presentation laptop and a cardioid lavalier microphone (Audio Technica AT898, 200 to 15,000 Hz response, 63 dB SNR) clipped to clothing near the mouth. These recordings were analyzed offline to transcribe patient responses, as well as to determine the time of articulatory onset and offset.

### SIGNAL ANALYSIS

Analyses were performed with trials time-locked to either picture presentation or to articulation onset. In all analyses, baseline was defined relative to the picture presentation (-500 to -100ms). Line noise was removed with zero-phase second-order Butterworth bandstop filters at 60, 120, and 180 Hz. Broadband high-gamma activity (BGA; 70 to 150 Hz) was extracted from the raw electrocorticography local field potential by a frequency domain bandpass Hilbert filter (paired sigmoid flanks with half-width 1.5 Hz). Power was then calculated as the squared envelope of the analytic signal, normalized as a percent of baseline activity. Power time series were downsampled to 200 Hz using a non-overlapping sliding window average.

### STRUCTURAL IMAGING

Preoperative anatomical MRI scans were obtained using a 3T whole-body MRI scanner (Philips Medical Systems) fitted with a 16-channel SENSE head coil. Images were collected using a magnetization-prepared 180° radiofrequency pulse and rapid gradient-echo sequence with 1-mm sagittal slices and an in-plane resolution of 0.938 × 0.938 mm. Pial surface reconstructions were computed with FreeSurfer (v5.1) (Dale et al., 1999) and imported to AFNI (Cox, 1996). Postoperative CT scans were registered to preoperative MRI scans for localization of electrodes relative to cortex. Grid electrode locations were determined by a recursive grid partitioning technique and then optimized using intraoperative photographs (Pieters et al., 2013). Depth electrode locations were informed by implantation trajectories from the Robotic Surgical Assistant (ROSA, Zimmer-Biomet) system.

### DELINEATION OF REGIONS OF INTEREST

Regions of interest were determined based on the Destrieux Atlas (Destrieux et al., 2010) and prior intracranial electrode studies (Forseth et al., 2021; Woolnough et al., 2022), resulting in an anatamo-functionally defined set of ROIs. The delineated ROIs rely on the Destrieux Atlas to parcellate the suprasylvian language production network. The following regions were defined from this atlas: inferior frontal sulcus (IFS), pars triangularis (parsTri), pars opercularis (parsOp), precentral sulcus (preCS), pre-and postcentral gyrus (preCG and postCG), subcentral gyrus (SCG), central sulcus (CS), Heschl’s gyrus (HG) and planum temporale (PT). Supplementary motor area (SMA), and preSMA were defined using geodesic radii around a defined center point, based on prior studies (Forseth et al., 2021; Woolnough et al., 2022). The boundary between SMA and preSMA was approximated with a vertical plane extending from the anterior commissure (Kim et al., 2010).

Electrodes within each region were identified based on their location on the population standard pial surface. Active electrodes were identified by evaluating BGA percent change from baseline in a peri-articulatory window from -500 to 500ms relative to articulation onset. If an electrode showed >20% mean BGA activation in this window, it was considered active. For analyses across ROIs, individual electrode responses were combined using hierarchical mixed effects models at each time point, nesting electrodes within patients.

### LINEAR MIXED EFFECTS MODELING

Linear mixed effects (LME) models were used to model BGA as a function of response time (RT) in each ROI. All modeling was performed in MATLAB using the fitlme() function. RT was modeled as a fixed effect, and electrode BGA as a random effect, nested within patient. Independent LMEs were fit at each time point in each ROI. Positive beta values indicate BGA increases with shorter RTs. Time points with significant betas were identified across the time course with a Benjamini-Hochberg FDR-corrected threshold at q = 0.05.

We also used LMEs in the coherent picture naming condition to model the relationship between BGA and four predictor variables: RT, number of syllables, phonological neighborhood, and word frequency, all derived from the Irvine Phonotactic Online Dictionary (Vaden et al., 2009).

## Results

### OVERVIEW

130 epilepsy patients completed a picture-naming task from the Snodgrass and Vanderwart (1980) picture set (Figure 1B) with sEEG (n = 86 implants) or SDE (n = 39 implants) recordings. Here, we focus on the control condition of the picture-naming task, in which subjects responded to incoherent images with the word ‘scrambled’ (Forseth et al., 2021) (Figure 1A). 66 ± 18 (mean ± SD) scrambled trials were presented and in total, the word ‘scrambled’ was produced 8,078 times across the cohort. We were interested in the neural correlates of the variability of this single stereotyped word across subjects. The bisyllabic word ‘scrambled’ begins with a complex consonant cluster [skL] (Figure 1F). The second syllable is often greatly reduced (Hayes, 2011), such that the spectral energy in the first syllable exceeds that of the second. In our dataset, we also observed variability in articulation of the final /d/. In effect, some subjects articulated ‘scrambled’ while others articulated ‘scramble’.

**Figure 1:**
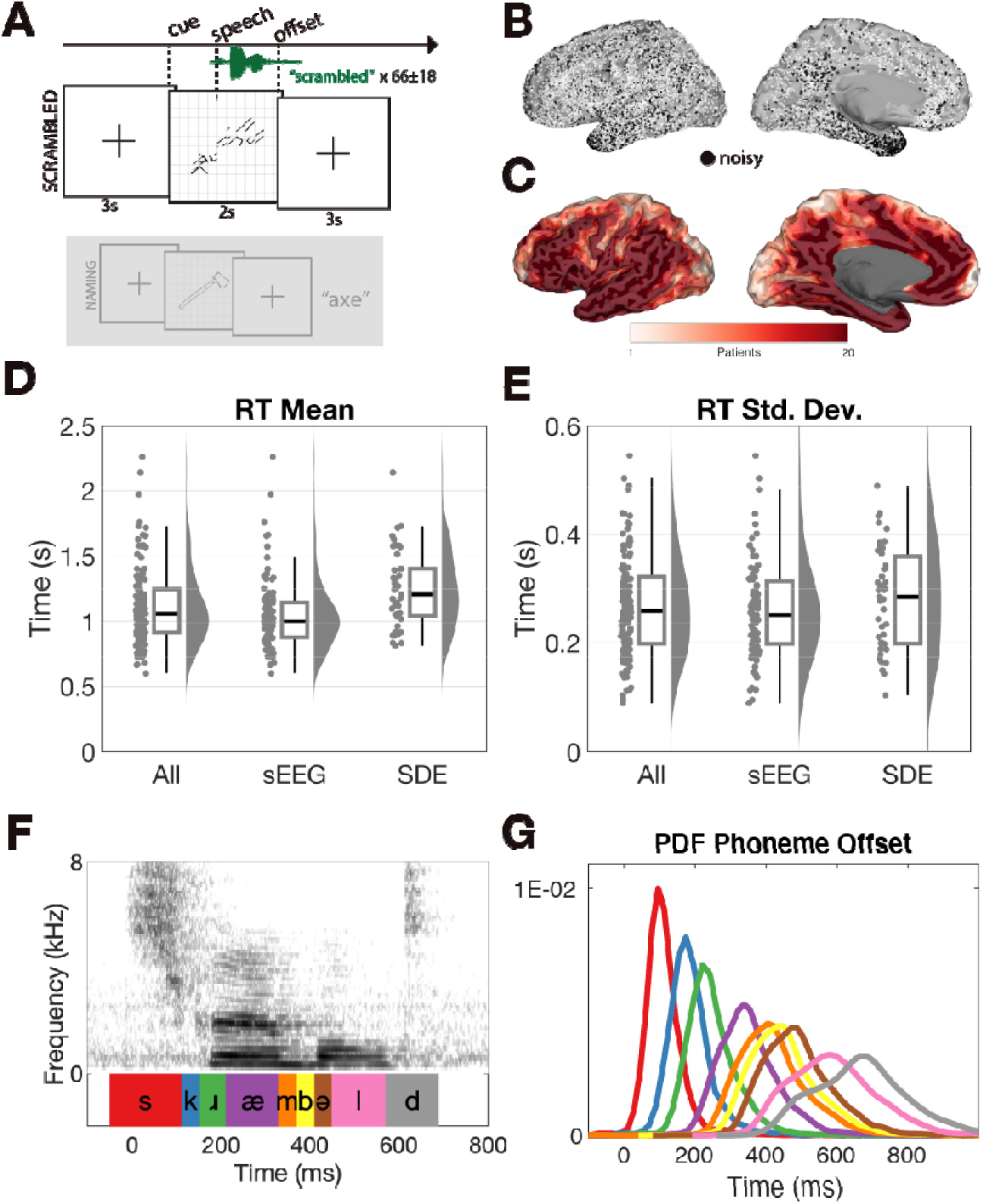
Experimental design, cortical coverage and response parameters. (A) Patients articulated ‘scrambled’ in response to an incoherent image in one third of trials (top) or named common objects in two-thirds of trials (bottom, gray overlay). (B) Coverage of language-dominant cortex with intracranial EEG. Top: Implanted electrodes registered to a standard semi-inflated pial surface. Electrodes included in analyses (white) or removed for epileptiform activity (black) are highlighted. (C) Representative coverage map across the cohort, with extensive coverage of both gyri and sulci. Mean (D) and standard deviation (E) of response time distributions for all patients and separately for SEEG implants only, and SDE only. Each data point represents the mean or standard deviation of RT for a single patient. (F) Spectrogram and phoneme boundaries identified by the Montreal Forced Aligner (McAuliffe et al., 2017) for one ‘scrambled’ production. Time zero is word onset as manually labeled by inspecting the spectrogram. (G) Probability distribution function of phoneme offsets color-coded from phonemes in (E). These distributions were generated from a subset of subjects (n = 40) with sufficiently clear audio for automated phonemic transcription.

### BEHAVIORAL ANALYSIS

Mean response time across the cohort was 1139 ± 298 ms. sEEG subjects responded, on average, 214 ms faster than SDE subjects (unpaired t-test, t(123) = 4.42, p < 0.001, 95% CI 118 to 310 ms), likely as patients with SDEs generally require a greater dosage of narcotics after surgery (Bernabei et al., 2021; Tandon et al., 2019), indexing the greater discomfort after a craniotomy. The mean duration of articulation of ‘scrambled’ was 638 ± 110 ms (Figure 1D). Using automated phonemic transcription, we identified approximate phoneme boundaries in subjects with sufficiently clear audio recordings (n = 40 patients) (Figure 1F). Articulatory variability of each phoneme in ‘scrambled’ accumulated variability of preceding phonemes; the first phoneme’s offset standard deviation was 59 ms relative to speech onset, while the last phoneme’s offset standard deviation was 131 ms relative to speech onset.

### 4D GLOBAL MEAN BROADBAND GAMMA POWER DYNAMICS

To inspect cortical activity across the cohort during stereotyped-word articulation, we utilized a surface-based mixed-effects multilevel analysis (SB-MEMA) technique which reliably co-registers activity across subjects onto a population standard surface, robustly accounts for trial and subject outliers, and infers activity at every point on the cortical surface (Conner et al., 2011; Kadipasaoglu et al., 2014). We used broadband gamma activity (BGA, 70 – 150 Hz) at each electrode to index local neural activity (Crone et al., 2006). Repeating SB-MEMA analyses in short, overlapping windows resulted in a 4D representation of cortical dynamics during the articulation of ‘scrambled’. We performed this analysis time-locked to speech onset (Video 1), and time-locked to stimulus onset (Video 2).

Around 200 ms after stimulus onset, SMA, precentral sulcus (preCS) and posterior inferior frontal sulcus (IFS; Video 2) became active. A short time later (∼250 ms), the anterior IFS and superior frontal sulcus became active. At 400 ms, premotor, primary motor, and sensorimotor regions became engaged and remained active for the duration of the articulation of the word. When visualizing timing of peak BGA activity, the SMA’s activity (Figure 2B, purple and white) is partially co-incident with IFS (purple) and precedes subsequent motor and premotor areas (orange).

**Figure 2:**
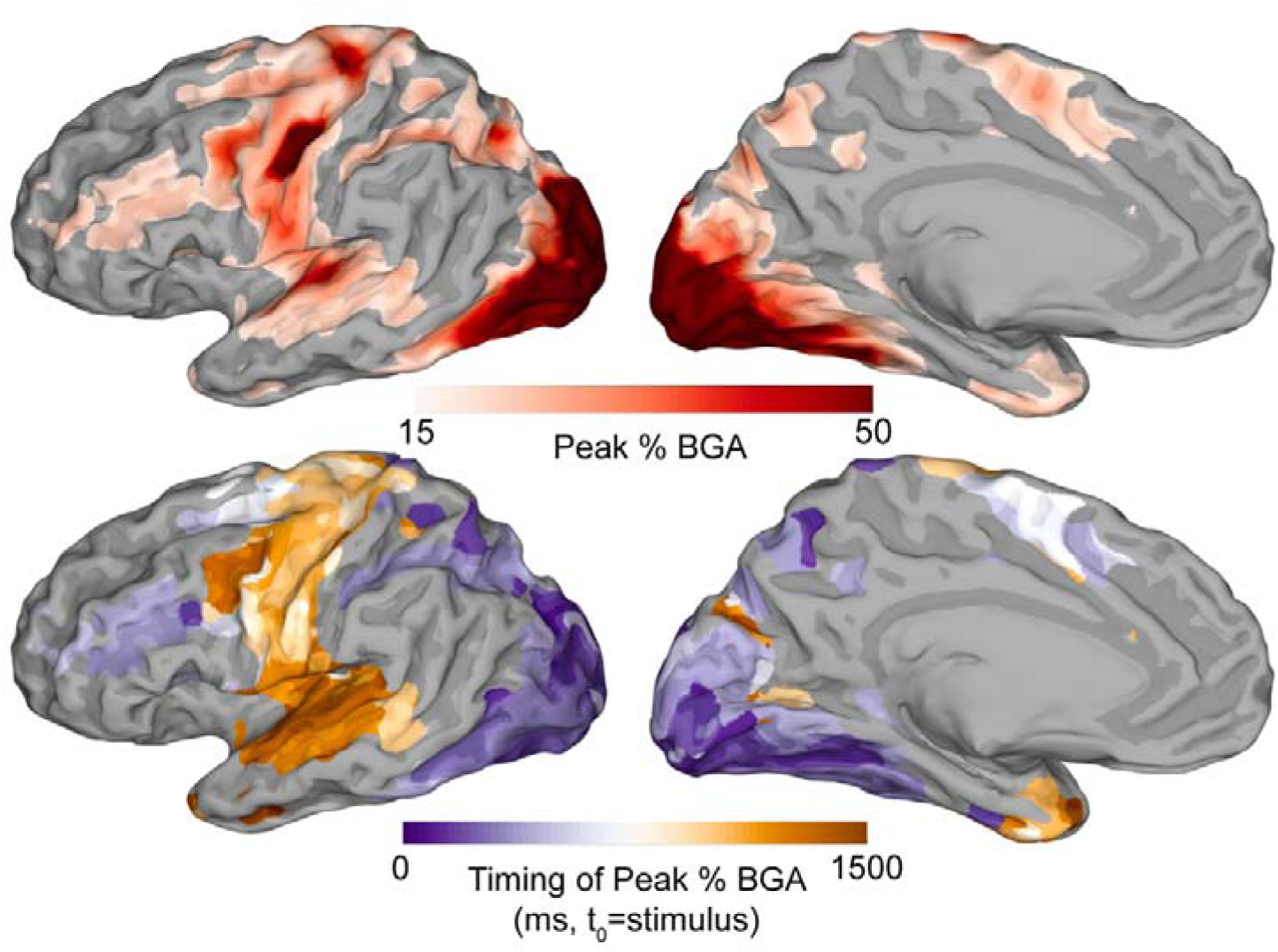
Cohort broadband gamma activity inferred with surface-based mixed-effects multilevel analysis. (A) Peak mean BGA percentage thresholded at 15% increase relative to baseline. (B) Timing of peak BGA. Refer to Video 1 for the full 4D rendition in both hemispheres.

Two functional zones, one in IFS and the other in preCS, became active 500 ms prior to speech onset (Video 1). Activity in both regions increased till 400 ms before speech onset after which IFS disengaged monotonically until articulation, when it became inactive, as predicted by prior work (Flinker et al., 2015). Meanwhile, preCS increased in activation and was involved throughout articulation.

We did not observe a somatosensory articulatory map of vSMC emerge at cohort level. In the population map, ventral and dorsal pre-and postcentral gyri engaged simultaneously. Given the existence of articulator maps in single-subject and few-subject studies (Bouchard et al., 2013), the absence of a population map is likely the result of inter-subject anatomical and functional variability. Indeed, at the single-subject level, BGA spread inferiorly and posteriorly in SCG from preCG to postCG at the time of speech onset (Video 3).

**Video 1: Cortex-wide 10-ms resolution broadband gamma activity inferred with surface-based mixed-effects multilevel analysis: articulation-locked.**

Time zero corresponds to articulation onset. The light grey distribution indicates cohort stimulus onset times. The dark grey distribution indicates cohort articulation offset times.

**Video 2: Broadband gamma activity inferred with surface-based mixed-effects multilevel analysis: stimulus-locked.**

**Video 3: Single-subject BGA across language-dominant hemisphere: articulation-locked.**

### BGA IN ARTICULATION REGIONS OF INTEREST

We next sought to compare SMA and preSMA activation time courses to that of other speech motor regions. Using the global activation maps (Figure 2; Video 1), we probed twelve anatomo-functional regions of interest (ROIs) in the language-dominant hemisphere (Figure 3; see Methods for ROI definitions). In addition to the supra-sylvian speech production network, we included Heschl’s gyrus and planum temporale given their proposed role in articulation monitoring and error detection (Forseth et al., 2020; Hickok, 2012). Activity in the SMA-proper peaked 300ms prior to articulation (∼600 ms post-stimulus) and declined rapidly at 100ms prior to articulation. PreSMA and SMA-proper were among the first regions activated and the first to deactivate (Figure 3B), starting around 500 ms after articulation onset, or just over 100 ms prior to median articulation offset (Figure 1F). Activity in precentral gyrus, subcentral gyrus, and postcentral gyrus peaked just after articulation onset, and these sites remained active even after articulation offset.

**Figure 3:**
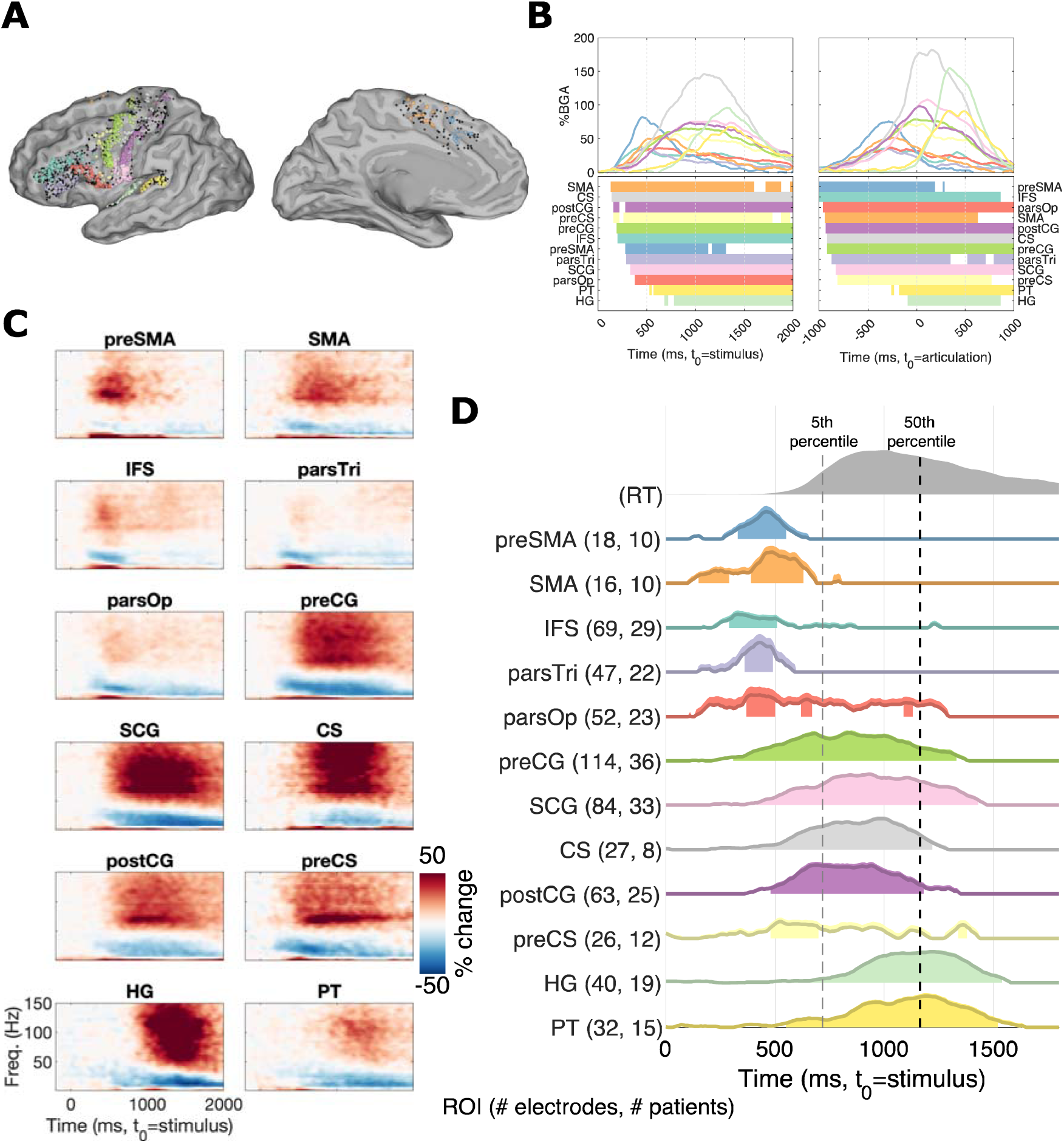
Language-dominant hemisphere broadband gamma activity in regions of interest: trial-averaged and single-trial response time correlation. (A) Regions of interest delineation on the standard cortical surface, colored according to region with inactive electrodes in black. (B) Average BGA time-locked to stimulus (left) and to speech onset (right) in the twelve ROIs. Periods of significant activation above baseline (FDR-corrected, q<0.01) are highlighted. (C) ROI-averaged spectrograms time-locked to stimulus onset in the twelve ROIs. (D) Linear mixed effects (LME) modeling of BGA as a function of RT in each ROI. LME beta values were half-wave rectified – interpretable as positive beta values indicating greater BGA in faster RT trials. The distribution of cohort RTs is plotted above in gray, with the dashed lines denoting fifth percentile and median RTs. Shaded area extending upwards from the beta curve denotes standard error. Time points which reach significance (FDR-corrected, q<0.05) are filled in under the curve. *Abbreviations*: IFS = inferior frontal sulcus, preCS = precentral sulcus, parsTri = pars triangularis, parsOp = pars opercularis, preSMA = pre-supplementary motor area, SMA = supplementary motor area-proper, preCG = precentral gyrus, SCG = subcentral gyrus, CS = central sulcus, postCG = postcentral gyrus, HG = Heschl’s gyrus, PT = planum temporale.

Spectrograms were calculated for each of the ROIs to inspect full spectrum changes relative to baseline (Figure 3C). In all ROIs, broadband gamma increases were expectedly accompanied by beta decreases. Pre-SMA beta decreases were smaller than other regions when scaled by BGA increases.

### SMA ACTIVITY PREDICTS TRIAL RESPONSE TIME

To further probe the role of SMA in speech initiation, we asked: how does single-trial BGA in SMA and preSMA contribute to prediction of RT? We ran linear mixed-effects models in each ROI, for each 10 ms interval, modeling BGA as a function of RT (fixed effect) and electrode, nested within subject (random effects); we found neural activity in SMA is modulated by RT early—just 270 ms after stimulus onset—and that preSMA and IFS followed SMA-proper about 50 ms later (Figure 3D). activity in pars opercularis, pars triangularis, and precentral sulcus did not co-vary with RT despite early activation (Figure 3D). Speech monitoring regions, Heschl’s gyrus and planum temporale, showed initial significance around the earliest articulation onsets given that they are time-locked to articulation.

### DRIVERS OF ACTIVATION IN SUPPLEMENTARY MOTOR AREA

The above analyses highlighted the early role of SMA-proper and preSMA in the intent to articulate. To investigate these areas further, and as we saw minimal distinctions between electrodes in preSMA and SMA-proper, we combined these ROIs into a single ROI to assess if these RT-driven electrodes are modulated by other behavioral correlates, e.g. articulatory complexity. To assess this, we tested the sensitivity of BGA in non-scrambled trials from the picture-naming task at the group level with linear mixed-effects modeling (Figure 4C). We were thus able to test how RT (relative to other behavioral predictors) contributes to BGA in a held-out dataset. The LME model (300 - 600 ms post-stimulus, r^2^ = 0.58) showed significant modulation by RT (t(5627) = -4.0, p < 0.001, 95% CI -35 to -12) and word frequency (t(5627) = -2.5, p = 0.012, 95% CI -29 to -4). Number of syllables (t(5627) = - 0.9, p = 0.35, 95% CI -16 to 6) and phonological neighborhood density (t(5627) = 1.1, p = 0.25, 95% CI -4 to -14) did not significantly modulate BGA in SMA.

**Figure 4:**
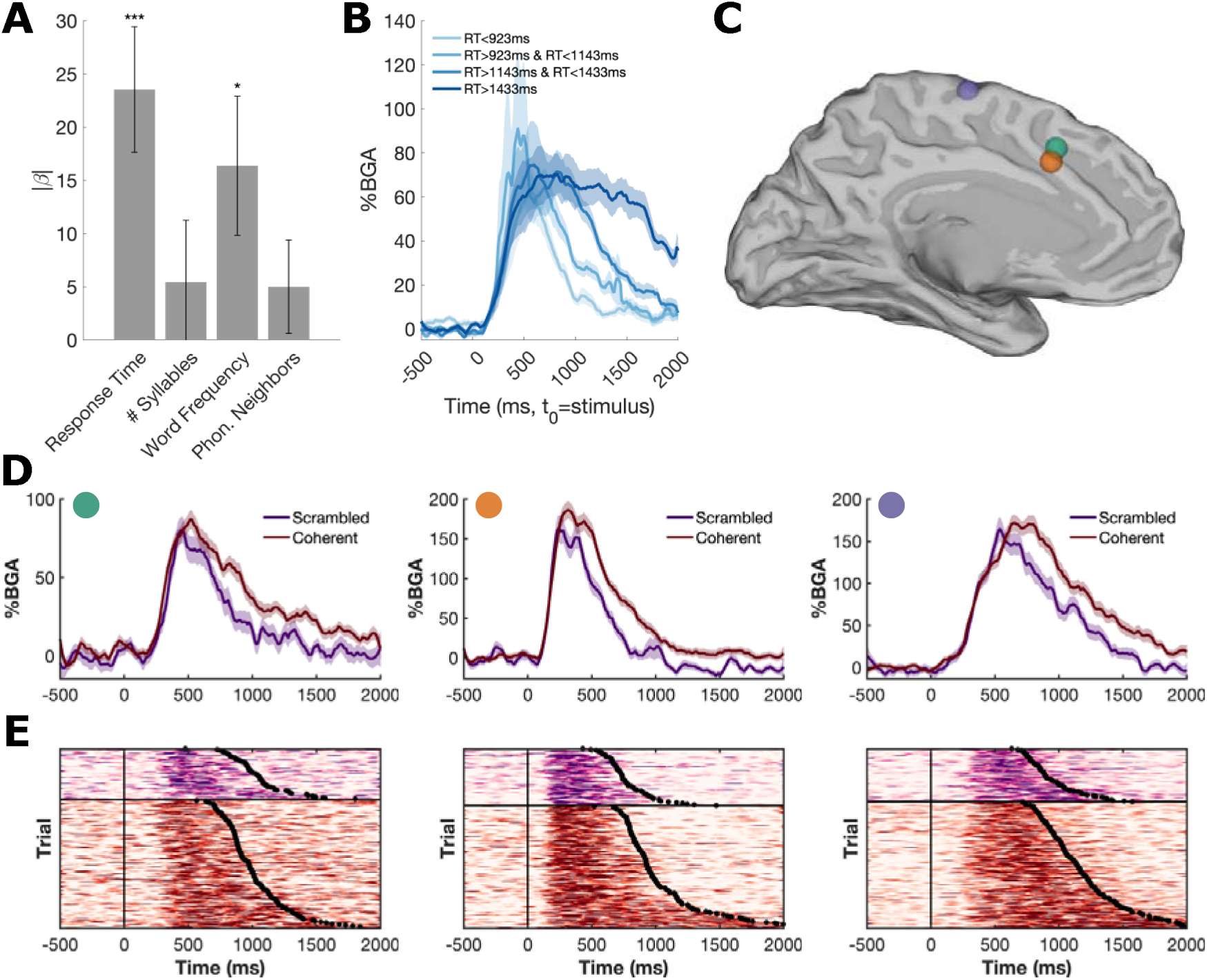
Functional broadband gamma activity in SMA for both scrambled and coherent picture naming conditions. (A) Linear mixed effects modeling of pre-articulatory BGA (300 – 600 ms post-stimulus onset) in the coherent naming trials as a function of four features: RT, number of syllables, word frequency, and phonological neighborhood. *** p < 0.001, * p < 0.05. (B) SMA BGA traces (mean ± SE) split by RT quartiles. (C,D,E) Exemplar electrodes from three different subjects: (C) locations in group-normalized space, (D) mean BGA traces for coherent (red) and scrambled (purple) trials and (E) trial-by-trial raster plots of BGA, sorted by trial RT.

To elaborate the stereotypical responses in SMA, we inspected activity at a single electrode level (Figure 4D,E). We found electrodes engaged at a predictable time after stimulus onset (200 - 300 ms in the exemplar electrodes, Figure 4D) and disengaged at a predictable time prior to speech onset. This pattern was only observable when electrodes were visualized in a trial-by-trial plot (Figure 4D, bottom) and not in the standard trial-averaged BGA trace (Figure 4D, top). This suggests that the activity at these electrodes lengthened with greater RT.

## Discussion

Speech initiation requires coordination across multiple cortical regions. The supplementary motor area, amongst other roles in speech (Hertrich et al., 2016), may relay a ‘go’ signal from the basal ganglia when high-level language areas and low-level sensorimotor areas are all ready to initiate a speech sequence (Guenther, 2016). Here, we analyzed intracranial recordings as subjects articulated a lexically-impoverished word ‘scrambled’ during a picture-naming task. We found evidence largely concordant with the idea that SMA is a key node in the healthy brain’s speech initiation network.

From a comprehensive 4-dimensional cortical atlas we extracted a cohort to show that the supplementary motor area along with inferior frontal sulcus are the earliest cortical regions to activate during cued, stereotyped word production (Figure 2; Figure 3B; Video 1, 2). Additionally, SMA activity was predictive of articulation onset at the single trial level and again it was one of the earliest regions to do so (Figure 3D). Finally, we uncovered a stereotypical BGA response in SMA during articulation preparation. SMA electrodes were modulated almost exclusively by response time, a conclusion supported by that fact that a) BGA was prolonged in trials with longer response time (Figure 4B–C) and b) other factors, like articulatory complexity, accounted for an insubstantial portion of BGA variance (Figure 4C). This is in line with the idea that SMA lies at the heart of the speech initiation circuit (Guenther, 2016). Our results do not directly support the idea that SMA initiates linguistic units smaller than words (like syllables or phonemes), nor the idea that SMA encodes indices of articulatory complexity like syllable count or phonological neighborhood; however, these factors may modulate SMA at smaller spatial scales than LFPs (Shima & Tanji, 2000).

Some electrodes in SMA showed remarkably consistent time-locking to intent-to-speak and articulation onset (Figure 4D). SMA activity *offset* coincided with vPMC and M1 activity *onset* at speech onset. This, along with existing literature, suggests that the SMA coordinates the concerted activation of vPMC and M1. But future research establishing a causal connection will need to elaborate on this speculation.

Previous non-invasive (Brendel et al., 2010) and invasive (IKEDA et al., 1992; Ohara et al., 2000) studies have yielded conflicting results regarding whether SMA is primarily involved in pre-movement planning window *prior* to speech onset or primarily engaged *during* speech execution, or both. Our results primarily support the preparation-window thesis; activity at the group (Figure 3B) and single-electrode level (Figure 4D-E) was constrained to the pre-speech window. There were, however, some electrodes which revealed execution-window (that is, *during* speech) activity.

### THE ROLE OF SMA IN SPEECH INITIATION AND TIMING

Penfield and Welch (1951) established a causal role of the SMA in speech production with local electrical cortical stimulation. Since then, the SMA has been further described as the site of ‘speech motor emission’ (Jonas, 1981) and of speech gesture sequencing (Ziegler et al., 1997). EEG studies revealed that SMA is the earliest region to peak in neural activity during speech production and that SMA activity peaks just prior to speech onset (Deecke et al., 1985; Grözinger et al., 1980). While the SMA is recognized in some current models of speech production (Guenther, 2016), it is absent in others (Hickok, 2012; Hickok et al., 2021). Recent evidence of SMA’s homologue in non-human primates has supported the idea that the SMA area mediates motor gesture release. Cadena-Valencia et al. (2018) showed that SMA maintains both internal (perceptual) and external (motor) rhythms in rhesus monkeys, suggesting that SMA is not only responsible for voluntary motor initiation, but also for managing rhythmic motor commands. Speech production, pseudo-rhythmic itself (Ghazanfar & Poeppel, 2014; ten Oever & Martin, 2021), may recruit SMA for monitoring and managing the release of speech motor commands at regular intervals (Hertrich et al., 2016). Our results corroborate the motor-release hypothesis for a single word but are agnostic to the question of phrase or discourse-level speech.

The DIVA/GODIVA model of speech production has developed to include the SMA (Bohland et al., 2010; Guenther, 2016). GODIVA hypothesizes that preSMA and IFS instantiate a “cognitive context” which holds context in memory until the appropriate motor gesture has been produced, at which point preSMA sends an initiation signal to SMA. SMA, in turn, interfaces with basal ganglia to “release” the planned speech acts (Guenther, 2016; Hertrich et al., 2016). Our results suggest that, at the LFP level, words are the units of release for the region, rather than the syllables or phonemes within the word; we observed activity prior to speech onset only, rather than activity prior syllables or phonemes (Figure 4). Our results begin to clarify the nature of SMA’s “release” of motor commands in the model. Previously, various possibilities existed for how this was achieved. For example, SMA could activate just prior to speech onset—independent of reaction time—if it were receiving a transient ‘go’ signal from the basal ganglia (Chang & Guenther, 2019). Our results, however, suggest that the stereotypical SMA response is time-locked to both intention to speak (cue onset) and the release of the motor command (speech onset). Thus, it is the *silencing* of BGA in SMA which indexes with the release of motor commands at the LFP scale.

SMA-proper and pre-SMA are known to have different cytoarchitectonic compositions (Ruan et al., 2018), functional connectivity (Ruan et al., 2018), and structural connectivity (Dick et al., 2019). Despite these differences, they often have similar activation patterns. Our results do not functionally separate these two regions.

### SMA IN PATHOLOGICAL SPEECH

The SMA has been implicated in a number of speech production disorders: namely, stuttering (Alm, 2004; Chang & Guenther, 2019; Chang & Zhu, 2013), apraxia of speech (Josephs et al., 2012; Utianski et al., 2018), SMA syndrome (Ardila, 2020; Pinson et al., 2021), and idiosyncratic aphasias (Berthier et al., 2015). SMA is often understood as a messenger node relaying sensorimotor context and timing information during production from basal ganglia and thalamus to appropriate cortical areas (Busan, 2020). Our results corroborate this hypothesis by providing evidence for the SMA’s role in speech initiation.

The high recovery rate of SMA syndrome (Nakajima et al., 2020; Pinson et al., 2021; Zentner et al., 1996) and SMA lesions (Ardila, 2020) suggests that while SMA is indispensable for speech initiation in a neurologically intact brain, the initiation network is degenerate (Sajid et al., 2020), with compensatory mechanisms readily available. The left and right SMAs may complement each other to a greater degree than other speech motor nodes, as SMA syndrome recovery is associated with increased right SMA connectivity with lateral speech regions (Pinson et al., 2021; Sailor et al., 2003).

### SMA IN BRAIN-COMPUTER INTERFACES

We suggest that future BCIs can leverage SMA to index timing of voluntary speech acts. SMA was the earliest predictor of speech onset. 250 ms after stimulus presentation, SMA activity significantly predicts response time. vSMC, which has been used to detect speech onset, becomes predictive more than 100 ms later, at 380 ms (Figure 2). In addition to offering an earlier prediction of voluntary speech acts than vSMC, SMA is a better speech detector in that a) it may be more active than vSMC during imagined, unexecuted speech (Park et al., 2015 Roland et al., 1980) and b) it houses single-site predictors of speech onset, which would alternatively have to be derived from the activity across many electrodes in vSMC (e.g., Moses et al., 2019).

### LIMITATIONS AND FUTURE DIRECTIONS

It is possible that the high-gamma LFP signal is not sufficiently resolved to expose syllabic or phonemic activity, and that smaller neural populations are tuned to these shorter motor commands. It is also worth considering that the BGA signal that we observed is likely not be speech-specific; pre-articulation BGA contribute to the *Bereitschaftspotential* (BP), or readiness potential (Deecke, 1987; Nachev et al., 2008) observed in human EEG prior to internally-generated movements, including speech (Grözinger et al., 1980; McArdle et al., 2009).

While the intracranial recordings in this study definitively show BGA in SMA during speech, we do not show that this set of SMA electrodes is speech-specific as we did not control for the initiation of non-speech motor acts.

Our results revealed a heterogeneous set of functional profiles at electrodes in the SMA. Denser coverage of the area will clarify how subregions of cortex pattern. In particular, we did not precisely delineate functional responses of SMA and preSMA. This may have been due to poorer sampling in preSMA specifically or across the medial wall more generally. On the other hand, some researchers have argued that SMA and preSMA do not form functionally discrete areas in the first place (Nachev et al., 2008).

Our results suggest that SMA is active primarily prior to articulation of a single word in a picture-naming task. Future studies should confirm whether the SMA is active primarily prior to phrase onset and before each word in a phrase. Our study concentrated on the most robust marker of single-unit activity (broadband gamma activity), but other frequency bands should be inspected. We used local field potentials in our analyses to show that the SMA is active only at the word level. It is possible that some sub-populations, or even single units, in SMA are active during word articulation and mediate the release of individual phonemes.

## Supporting information

Video 1 scrambled MEMA articulation

Video 2 scrambled MEMA stimulus

Video 3 TA715 articulation

## Acknowledgements

We express our gratitude to all the patients who participated in this study; the neurologists at the Texas Comprehensive Epilepsy Program who participated in the care of these patients; and the nurses and technicians in the Epilepsy Monitoring Unit at Memorial Hermann Hospital who helped make this research possible. This work was supported by NIH R01 DC014589 and the University of Texas System funding for the Texas Institute for Restorative Neurotechnologies.

## Author Contributions

Conceptualization: LB, KJF and NT; Data Curation: KJF and PSR; Formal Analysis and Visualization: LB and OW; Writing - Original Draft: LB; Writing - Review and Editing: LB, KJF, OW and NT; Funding Acquisition: NT; Neurosurgical Procedures: NT.

## References

1. Alario, F.-X., Chainay, H., Lehericy, S., & Cohen, L. (2006). The role of the supplementary motor area (SMA) in word production. Brain Research, 1076(1), 129–143. https://doi.org/10.1016/j.brainres.2005.11.104

2. Alm, P. A. (2004). Stuttering and the basal ganglia circuits: a critical review of possible relations. Journal of Communication Disorders, 37(4), 325–369. https://doi.org/10.1016/j.jcomdis.2004.03.001

3. Ardila, A. (2020). Supplementary motor area aphasia revisited. Journal of Neurolinguistics, 54, 100888. https://doi.org/10.1016/j.jneuroling.2020.100888

4. Bernabei, J. M., Arnold, T. C., Shah, P., Revell, A., Ong, I. Z., Kini, L. G., Stein, J. M., Shinohara, R. T., Lucas, T. H., Davis, K. A., Bassett, D. S., & Litt, B. (2021). Electrocorticography and stereo EEG provide distinct measures of brain connectivity: implications for network models. Brain Communications, 3(3), fcab156. https://doi.org/10.1093/braincomms/fcab156

5. Berthier, M., Dávila, G., Moreno-Torres, I., Beltrán-Corbellini, Á., Santana-Moreno, D., Torres-Prioris, M. J., Massone, M. I., Cruces, R., Roé Vellvé, N., & Thurnhofer-Hemsi, K. (2015). Loss of regional accent after damage to the speech production network. Frontiers in Human Neuroscience, 9. https://www.frontiersin.org/article/10.3389/fnhum.2015.00610

6. Bohland, J. W., Bullock, D., & Guenther, F. H. (2010). Neural representations and mechanisms for the performance of simple speech sequences. Journal of Cognitive Neuroscience, 22(7), 1504–1529. https://doi.org/10.1162/jocn.2009.21306

7. Bohland, J. W., & Guenther, F. H. (2006). An fMRI investigation of syllable sequence production. NeuroImage, 32(2), 821–841. https://doi.org/10.1016/j.neuroimage.2006.04.173

8. Bouchard, K. E., Mesgarani, N., Johnson, K., & Chang, E. F. (2013). Functional organization of human sensorimotor cortex for speech articulation. Nature, 495(7441), 327–332. https://doi.org/10.1038/nature11911

9. Brendel, B., Hertrich, I., Erb, M., Lindner, A., Riecker, A., Grodd, W., & Ackermann, H. (2010). The contribution of mesiofrontal cortex to the preparation and execution of repetitive syllable productions: An fMRI study. NeuroImage, 50(3), 1219–1230. https://doi.org/10.1016/j.neuroimage.2010.01.039

10. Busan, P. (2020). Developmental stuttering and the role of the supplementary motor cortex. Journal of Fluency Disorders, 64, 105763. https://doi.org/10.1016/j.jfludis.2020.105763

11. Cadena-Valencia, J., García-Garibay, O., Merchant, H., Jazayeri, M., & Lafuente, V. de. (2018, October 22). Entrainment and maintenance of an internal metronome in supplementary motor area. ELife; eLife Sciences Publications Limited. https://doi.org/10.7554/eLife.38983

12. Castellucci, G. A., Kovach, C. K., Howard, M. A., Greenlee, J. D. W., & Long, M. A. (2022). A speech planning network for interactive language use. Nature, 1–6. https://doi.org/10.1038/s41586-021-04270-z

13. Catani, M., Dell’Acqua, F., Vergani, F., Malik, F., Hodge, H., Roy, P., Valabregue, R., & Thiebaut de Schotten, M. (2012). Short frontal lobe connections of the human brain. Cortex, 48(2), 273–291. https://doi.org/10.1016/j.cortex.2011.12.001

14. Chang, S.-E., & Guenther, F. H. (2019). Involvement of the Cortico-Basal Ganglia-Thalamocortical Loop in Developmental Stuttering. Frontiers in Psychology, 10, 3088. https://doi.org/10.3389/fpsyg.2019.03088

15. Chang, S.-E., & Zhu, D. C. (2013). Neural network connectivity differences in children who stutter. Brain, 136(12), 3709–3726. https://doi.org/10.1093/brain/awt275

16. Chartier, J., Anumanchipalli, G. K., Johnson, K., & Chang, E. F. (2018). Encoding of Articulatory Kinematic Trajectories in Human Speech Sensorimotor Cortex. Neuron, 98(5), 1042–1054.e4. https://doi.org/10.1016/j.neuron.2018.04.031

17. Conant, D., Bouchard, K. E., & Chang, E. F. (2014). Speech map in the human ventral sensory-motor cortex. Current Opinion in Neurobiology, 24, 63–67. https://doi.org/10.1016/j.conb.2013.08.015

18. Conner, C. R., Chen, G., Pieters, T. A., & Tandon, N. (2014). Category Specific Spatial Dissociations of Parallel Processes Underlying Visual Naming. Cerebral Cortex, 24(10), 2741–2750. https://doi.org/10.1093/cercor/bht130

19. Conner, C. R., Ellmore, T. M., Pieters, T. A., DiSano, M. A., & Tandon, N. (2011). Variability of the Relationship between Electrophysiology and BOLD-fMRI across Cortical Regions in Humans. The Journal of Neuroscience, 31(36), 12855–12865. https://doi.org/10.1523/JNEUROSCI.1457-11.2011

20. Conner, C. R., Kadipasaoglu, C. M., Shouval, H. Z., Hickok, G., & Tandon, N. (2019). Network dynamics of Broca’s area during word selection. PLOS ONE, 14(12), e0225756. https://doi.org/10.1371/journal.pone.0225756

21. Cox, R. W. (1996). AFNI: software for analysis and visualization of functional magnetic resonance neuroimages. Computers and Biomedical Research, an International Journal, 29(3), 162–173. https://doi.org/10.1006/cbmr.1996.0014

22. Crone, N. E., Sinai, A., & Korzeniewska, A. (2006). High-frequency gamma oscillations and human brain mapping with electrocorticography. Progress in Brain Research, 159, 275–295. https://doi.org/10.1016/S0079-6123(06)59019-3

23. Dale, A. M., Fischl, B., & Sereno, M. I. (1999). Cortical surface-based analysis. I. Segmentation and surface reconstruction. NeuroImage, 9(2), 179–194. https://doi.org/10.1006/nimg.1998.0395

24. Deecke, L. (1987). Bereitschaftspotential as an indicator of movement preparation in supplementary motor area and motor cortex. Ciba Foundation Symposium, 132, 231–250. https://doi.org/10.1002/9780470513545.ch14

25. Deecke, L., Kornhuber, H. H., Lang, W., Lang, M., & Schreiber, H. (1985). Timing function of the frontal cortex in sequential motor and learning tasks. Human Neurobiology, 4(3), 143–154.

26. Destrieux, C., FISCHL, B., DALE, A., & HALGREN, E. (2010). Automatic parcellation of human cortical gyri and sulci using standard anatomical nomenclature. NeuroImage, 53(1), 1–15. https://doi.org/10.1016/j.neuroimage.2010.06.010

27. Dick, A. S., Garic, D., Graziano, P., & Tremblay, P. (2019). The frontal aslant tract (FAT) and its role in speech, language and executive function. Cortex, 111, 148–163. https://doi.org/10.1016/j.cortex.2018.10.015

28. Flinker, A., Korzeniewska, A., Shestyuk, A. Y., Franaszczuk, P. J., Dronkers, N. F., Knight, R. T., & Crone, N. E. (2015). Redefining the role of Broca’s area in speech. Proceedings of the National Academy of Sciences, 112(9), 2871–2875. https://doi.org/10.1073/pnas.1414491112

29. Forseth, K. J., Hickok, G., Rollo, P. S., & Tandon, N. (2020). Language prediction mechanisms in human auditory cortex. Nature Communications, 11(1), 5240. https://doi.org/10.1038/s41467-020-19010-6

30. Forseth, K. J., Kadipasaoglu, C. M., Conner, C. R., Hickok, G., Knight, R. T., & Tandon, N. (2018). A lexical semantic hub for heteromodal naming in middle fusiform gyrus. Brain, 141(7), 2112–2126. https://doi.org/10.1093/brain/awy120

31. Forseth, K. J., Pitkow, X., Fischer-Baum, S., & Tandon, N. (2021). What The Brain Does As We Speak (p. 2021.02.05.429841). bioRxiv. https://doi.org/10.1101/2021.02.05.429841

32. Ghazanfar, A. A., & Poeppel, D. (2014). The neurophysiology and evolution of the speech rhythm. In The cognitive neurosciences, 5th ed (pp. 629–638). MIT Press.

33. Goldenholz, D. M., Ahlfors, S. P., Hämäläinen, M. S., Sharon, D., Ishitobi, M., Vaina, L. M., & Stufflebeam, S. M. (2008). Mapping the signal-to-noise-ratios of cortical sources in magnetoencephalography and electroencephalography. Human Brain Mapping, 30(4), 1077–1086. https://doi.org/10.1002/hbm.20571

34. González-Martínez, J., Bulacio, J., Thompson, S., Gale, J., Smithason, S., Najm, I., & Bingaman, W. (2016). Technique, Results, and Complications Related to Robot-Assisted Stereoelectroencephalography. Neurosurgery, 78(2), 169–180. https://doi.org/10.1227/NEU.0000000000001034

35. Gonzalez-Martinez, J., Mullin, J., Vadera, S., Bulacio, J., Hughes, G., Jones, S., Enatsu, R., & Najm, I. (2014). Stereotactic placement of depth electrodes in medically intractable epilepsy: Technical note. Journal of Neurosurgery, 120(3), 639–644. https://doi.org/10.3171/2013.11.JNS13635

36. Grözinger, B., Kornhuber, H. H., Kriebel, J., Szirtes, J., & Westphal, K. T. P. (1980). The Bereitschaftspotential Preceding the Act of Speaking. Also an Analysis of Artifacts. In

37. H. H. Kornhubek & L. Deecke (Eds.), Progress in Brain Research (Vol. 54, pp. 798– 804). Elsevier. https://doi.org/10.1016/S0079-6123(08)61705-7

38. Guenther, F. H. (2016). Neural Control of Speech. MIT Press. Hayes, B. (2011). Introductory Phonology. John Wiley & Sons.

39. Hertrich, I., Dietrich, S., & Ackermann, H. (2016). The role of the supplementary motor area for speech and language processing. Neuroscience & Biobehavioral Reviews, 68, 602–610. https://doi.org/10.1016/j.neubiorev.2016.06.030

40. Hickok, G. (2012). Computational neuroanatomy of speech production. Nature Reviews Neuroscience, 13(2), 135–145. https://doi.org/10.1038/nrn3158

41. Hickok, G., Venezia, J. H., & Teghipco, A. (2021). Beyond Broca: Neural Architecture and Evolution of a Dual Motor Speech Coordination System [Preprint]. PsyArXiv. https://doi.org/10.31234/osf.io/tewna

42. IKEDA, A., LÜDERS, H. O., BURGESS, R. C.,& SHIBASAKI, H. (1992). MOVEMENT-RELATED POTENTIALS RECORDED FROM SUPPLEMENTARY MOTOR AREA AND PRIMARY MOTOR AREA: ROLE OF SUPPLEMENTARY MOTOR AREA IN VOLUNTARY MOVEMENTS. Brain, 115(4), 1017–1043. https://doi.org/10.1093/brain/115.4.1017

43. Jonas, S. (1981). The supplementary motor region and speech emission. Journal of Communication Disorders, 14(5), 349–373. https://doi.org/10.1016/0021-9924(81)90019-8

44. Josephs, K. A., Duffy, J. R., Strand, E. A., Machulda, M. M., Senjem, M. L., Master, A. V., Lowe, V. J., Jack, C. R., & Whitwell, J. L. (2012). Characterizing a neurodegenerative syndrome: primary progressive apraxia of speech. Brain: A Journal of Neurology, 135(Pt 5), 1522–1536. https://doi.org/10.1093/brain/aws032

45. Kadipasaoglu, C. M., Baboyan, V. G., Conner, C. R., Chen, G., Saad, Z. S., & Tandon, N. (2014). Surface-based mixed effects multilevel analysis of grouped human electrocorticography. NeuroImage, 101, 215–224. https://doi.org/10.1016/j.neuroimage.2014.07.006

46. Kermadi, Y. Liu, A. Tempini E.M. Ro, I. (1997). Effects of reversible inactivation of the supplementary motor area (SMA) on unimanual grasp and bimanual pull and grasp performance in monkeys. Somatosensory & Motor Research, 14(4), 268–280. https://doi.org/10.1080/08990229770980

47. Kim, J.-H., Lee, J.-M., Jo, H. J., Kim, S. H., Lee, J. H., Kim, S. T., Seo, S. W., Cox, R. W., Na, D. L., Kim, S. I., & Saad, Z. S. (2010). Defining functional SMA and pre-SMA subregions in human MFC using resting state fMRI: functional connectivity-based parcellation method. NeuroImage, 49(3), 2375. https://doi.org/10.1016/j.neuroimage.2009.10.016

48. Krainik, A., Lehéricy, S., Duffau, H., Capelle, L., Chainay, H., Cornu, P., Cohen, L., Boch, A.-L., Mangin, J.-F., Le Bihan, D., & Marsault, C. (2003). Postoperative speech disorder after medial frontal surgery: role of the supplementary motor area. Neurology, 60(4), 587–594. https://doi.org/10.1212/01.wnl.0000048206.07837.59

49. Laplane, D., Talairach, J., Meininger, V., Bancaud, J., & Orgogozo, J. M. (1977). Clinical consequences of corticectomies involving the supplementary motor area in man. Journal of the Neurological Sciences, 34(3), 301–314. https://doi.org/10.1016/0022-510x(77)90148-4

50. Lu, J., Zhao, Z., Zhang, J., Wu, B., Zhu, Y., Chang, E. F., Wu, J., Duffau, H., & Berger, M. S. (2021). Functional maps of direct electrical stimulation-induced speech arrest and anomia: a multicentre retrospective study. Brain, 144(8), 2541–2553. https://doi.org/10.1093/brain/awab125

51. McArdle, J. J., Mari, Z., Pursley, R. H., Schulz, G. M., & Braun, A. R. (2009). Electrophysiological evidence of functional integration between the language and motor systems in the brain: A study of the speech Bereitschaftspotential. Clinical Neurophysiology_: Official Journal of the International Federation of Clinical Neurophysiology, 120(2), 275–284. https://doi.org/10.1016/j.clinph.2008.10.159

52. Moses, D. A., Leonard, M. K., Makin, J. G., & Chang, E. F. (2019). Real-time decoding of question-and-answer speech dialogue using human cortical activity. Nature Communications, 10(1), 3096. https://doi.org/10.1038/s41467-019-10994-4

53. Mugler, E. M., Tate, M. C., Livescu, K., Templer, J. W., Goldrick, M. A., & Slutzky, M. W. (2018). Differential Representation of Articulatory Gestures and Phonemes in Precentral and Inferior Frontal Gyri. Journal of Neuroscience, 38(46), 9803–9813. https://doi.org/10.1523/JNEUROSCI.1206-18.2018

54. Nachev, P., Kennard, C., & Husain, M. (2008). Functional role of the supplementary and pre-supplementary motor areas. Nature Reviews Neuroscience, 9(11), 856–869. https://doi.org/10.1038/nrn2478

55. Nakajima, R., Kinoshita, M., Yahata, T., & Nakada, M. (2020). Recovery time from supplementary motor area syndrome: relationship to postoperative day 7 paralysis and damage of the cingulum. Journal of Neurosurgery, 132(3), 865–874. https://doi.org/10.3171/2018.10.JNS182391

56. Ohara, S., Ikeda, A., Kunieda, T., Yazawa, S., Baba, K., Nagamine, T., Taki, W., Hashimoto, N., Mihara, T., & Shibasaki, H. (2000). Movement-related change of electrocorticographic activity in human supplementary motor area proper. Brain, 123(6), 1203–1215. https://doi.org/10.1093/brain/123.6.1203

57. Park, C., Chang, W. H., Lee, M., Kwon, G. H., Kim, L., Kim, S. T., & Kim, Y.-H. (2015). Which motor cortical region best predicts imagined movement? NeuroImage, 113, 101–110. https://doi.org/10.1016/j.neuroimage.2015.03.033

58. Penfield, W., & Welch, K. (1951). The supplementary motor area of the cerebral cortex; a clinical and experimental study. A.M.A. Archives of Neurology and Psychiatry, 66(3), 289–317. https://doi.org/10.1001/archneurpsyc.1951.02320090038004

59. Pieters, T. A., Conner, C. R., & Tandon, N. (2013). Recursive grid partitioning on a cortical surface model: an optimized technique for the localization of implanted subdural electrodes. Journal of Neurosurgery, 118(5), 1086–1097. https://doi.org/10.3171/2013.2.JNS121450

60. Pinson, H., Van Lerbeirghe, J., Vanhauwaert, D., Van Damme, O., Hallaert, G., & Kalala, J.-P. (2021). The supplementary motor area syndrome: a neurosurgical review. Neurosurgical Review. https://doi.org/10.1007/s10143-021-01566-6

61. Roland, P. E., Larsen, B., Lassen, N. A., & Skinhøj, E. (1980). Supplementary motor area and other cortical areas in organization of voluntary movements in man. Journal of Neurophysiology, 43(1), 118–136. https://doi.org/10.1152/jn.1980.43.1.118

62. Rollo, P. S., Rollo, M. J., Zhu, P., Woolnough, O., & Tandon, N. (2020). Oblique trajectory angles in robotic stereo-electroencephalography. Journal of Neurosurgery, 135(1), 245–254. https://doi.org/10.3171/2020.5.JNS20975

63. Ruan, J., Bludau, S., Palomero-Gallagher, N., Caspers, S., Mohlberg, H., Eickhoff, S. B., Seitz, R. J., & Amunts, K. (2018). Cytoarchitecture, probability maps, and functions of the human supplementary and pre-supplementary motor areas. Brain Structure and Function, 223(9), 4169–4186. https://doi.org/10.1007/s00429-018-1738-6

64. Sailor, J., Meyerand, M. E., Moritz, C. H., Fine, J., Nelson, L., Badie, B., & Haughton, V. M. (2003). Supplementary Motor Area Activation in Patients with Frontal Lobe Tumors and Arteriovenous Malformations. 6.

65. Sajid, N., Parr, T., Hope, T. M., Price, C. J., & Friston, K. J. (2020). Degeneracy and Redundancy in Active Inference. Cerebral Cortex, 30(11), 5750–5766. https://doi.org/10.1093/cercor/bhaa148

66. Shima, K., & Tanji, J. (2000). Neuronal Activity in the Supplementary and Presupplementary Motor Areas for Temporal Organization of Multiple Movements. Journal of Neurophysiology, 84(4), 2148–2160. https://doi.org/10.1152/jn.2000.84.4.2148

67. Snodgrass, J. G., & Vanderwart, M. (1980). A standardized set of 260 pictures: Norms for name agreement, image agreement, familiarity, and visual complexity. Journal of Experimental Psychology: Human Learning and Memory, 6(2), 174–215. https://doi.org/10.1037/0278-7393.6.2.174

68. Tandon, N., & Luders, H. (2008). Cortical mapping by electrical stimulation of subdural electrodes: language areas. In Textbook of Epilepsy Surgery (pp. 1001–1015). McGraw Hill.

69. Tandon, N., Tong, B. A., Friedman, E. R., Johnson, J. A., Von Allmen, G., Thomas, M. S., Hope, O. A., Kalamangalam, G. P., Slater, J. D., & Thompson, S. A. (2019). Analysis of Morbidity and Outcomes Associated With Use of Subdural Grids vs Stereoelectroencephalography in Patients With Intractable Epilepsy. JAMA Neurology, 76(6), 672–681. https://doi.org/10.1001/jamaneurol.2019.0098

70. Tanji, J., & Shima, K. (1994). Role for supplementary motor area cells in planning several movements ahead. Nature, 371(6496), 413–416. https://doi.org/10.1038/371413a0

71. ten Oever, S., & Martin, A. E. (2021). An oscillating computational model can track pseudo-rhythmic speech by using linguistic predictions. ELife, 10, e68066. https://doi.org/10.7554/eLife.68066

72. Utianski, R. L., Duffy, J. R., Clark, H. M., Strand, E. A., Botha, H., Schwarz, C. G., Machulda, M. M., Senjem, M. L., Spychalla, A. J., Jack, C. R., Petersen, R. C., Lowe, V. J., Whitwell, J. L., & Josephs, K. A. (2018). Prosodic and Phonetic Subtypes of Primary Progressive Apraxia of Speech. Brain and Language, 184, 54–65. https://doi.org/10.1016/j.bandl.2018.06.004

73. Vaden, K. I., Halpin, H. R., & Hickok, G. (2009). Irvine Phonotactic Online Dictionary, Version 2.0.

74. Woolnough, O., Donos, C., Curtis, A., Rollo, P. S., Roccaforte, Z. J., Dehaene, S., Fischer-Baum, S., & Tandon, N. (2022). A Spatiotemporal Map of Reading Aloud. Journal of Neuroscience, 42(27), 5438–5450. https://doi.org/10.1523/JNEUROSCI.2324-21.2022

75. Woolnough, O., Forseth, K. J., Rollo, P. S., & Tandon, N. (2019). Uncovering the functional anatomy of the human insula during speech. ELife, 8, e53086. https://doi.org/10.7554/eLife.53086

76. Zentner, J., Hufnagel, A., Pechstein, U., Wolf, H. K., & Schramm, J. (1996). Functional results after resective procedures involving the supplementary motor area. Journal of Neurosurgery, 85(4), 542–549. https://doi.org/10.3171/jns.1996.85.4.0542

77. Ziegler, W., Kilian, B., & Deger, K. (1997). The role of the left mesial frontal cortex in fluent speech: Evidence from a case of left supplementary motor area hemorrhage. Neuropsychologia, 35(9), 1197–1208. https://doi.org/10.1016/S0028-3932(97)00040-7

